# Evaluation of scalp-based targeting methods of dorsolateral prefrontal cortex for transcranial magnetic stimulation therapy: Electroencephalogram cap, Beam F3, CPC F3, and adjusted Beam F3

**DOI:** 10.1101/2024.09.28.615621

**Authors:** Yihan Jiang, Yuanyuan Chen, Lijiang Wei, Farui Liu, Zeqing Zheng, Zong Zhang, Zheng Li, Yingying Tang, Jijun Wang, Qing Xie, Chuanxin M. Niu, Chaozhe Zhu

**Author notes:** Address correspondence to: Chaozhe Zhu, Ph.D. State Key Lab of Cognitive Neuroscience and Learning & IDG/McGovern Institute for Brain Research, Beijing Normal University, Beijing 100875, China., Chuanxin M. Niu, Ph.D., Department of Rehabilitation Medicine, Shanghai Jiao Tong University School of Medicine, Shanghai 200025, China. These authors contributed equally to this work (Yihan Jiang, Yuanyuan Chen).

## Abstract

**Background:** Repetitive transcranial magnetic stimulation targeting the left dorsolateral prefrontal cortex (DLPFC) offers a promising approach for treating depression. Scalp-based targeting methods, including EEG Cap, Beam F3, CPC F3, and adjusted Beam F3, have evolved to improve clinical application.

**Obejectives:** To evaluate and compare the accuracy, reliability, speed, and training efficacy of these four methods.

**Methods:** Three trained technicians conducted repeated DLPFC targeting measurements on 10 3D-printed head models. Accuracy was assessed by calculating targeting error and electric field strength at the subgenual anterior cingulate cortex anti-correlated peak. Reliability was evaluated through inter- and intra-rater variability, and training efficacy was examined by comparing targeting errors between trained technicians and novices.

**Results:** CPC F3, Beam F3, and adjusted Beam F3 demonstrated comparable accuracy in terms of electric field strength, but CPC F3 had the lowest targeting error (4.2 mm). CPC F3 also exhibited the highest reliability (inter-rater variability at 3.3 mm, intra-rater variability at 2.3 mm). The EEG cap was the fastest method (80 seconds), while CPC F3 was faster than Beam F3 but slower than EEG Cap. No significant differences in targeting error between trained technicians and novices were noted with CPC F3 and adjusted Beam F3.

**Conclusion:** The comparative analysis of four scalp-based targeting methods provides insights that can enhance clinical decision-making and the practical application. While accuracy among CPC F3, Beam F3, and adjusted Beam F3 was comparable, CPC F3 emerges as highly practical due to its significantly better reliability, training efficacy, moderate speed, and straightforward implementation.

## Introduction

Depression poses a profound challenge within global mental health^1^. Across super-regions, at least approximately 2% of the population grapples with severe depression^2^, many of whom may require treatment. Repetitive transcranial magnetic stimulation (rTMS) has proven effective in treating depression. Due to its non-invasive and minor side effects, rTMS occupies a distinct position within the array of therapeutic options for depression. Commonly, high-frequency stimulation over the left dorsolateral prefrontal cortex (DLPFC) is employed as a standard rTMS treatment protocol^3^. A crucial element of this protocol is the precise placement of the coil to target the DLPFC.

Scalp-based targeting methods facilitate the measurement of DLPFC for coil placement in depression treatment. By avoiding the expensive need for neural navigation^4^ and magnetic resonance imaging (MRI) scans^5, 6^, these methods potentially reducing the overall cost of medical treatments for patients^7^. Additionally, their straightforward implementation not only shortens treatment durations but also simplifies the training process for novices. Scalp-based targeting methods have been evolving to enhance clinical utilization. Initially, the coil placement for rTMS in depression treatment was approximately 5 cm anterior to the motor cortex hand area^8^. However, this placement omits variations in head size, leading to insufficient DLPFC targeting in more than one-third of cases^9, 10^. The F3 position, measured based on the percentages of the head circumference and distances between four key anatomical landmarks, better accommodates different head sizes. Manually assessing F3, as defined by the original International 10-20 System, involves identifying 14 markers^11^, a process that is both time-consuming and prone to introducing cumulative errors. Therefore, a specialized electroencephalogram (EEG) cap might be preferable. Alternatively, the Beam F3 method involves scalp measurements guided by accompanying software or a website^12^. This method effectively resolves the challenges arising from the cost and limited availability of EEG caps. More recently, the continuous proportional coordinate (CPC) system has been proposed to describe any scalp point using a pair of proportional coordinates^13^. Using a structural MRI database of 114 persons, large-sample-based average coordinates for F3, CPC F3 (0.27,0.34), were established^14^. Additionally, research by Fox et al. found the negative functional connectivity between the DLPFC and the subgenual anterior cingulate cortex (sgACC) serves as a predictive marker for TMS response in depression, highlighting the importance of targeting based on functional connectivity^15^. Building on this, an adjusted version of the Beam F3 method has been proposed to target the coordinate most strongly anti-correlated with the sgACC in the DLPFC at the group level^16^.

With the development of scalp-based targeting methods, the investigation of these approaches by researchers has increased. Nikolin^17^ and Trapp^18^ et al. have assessed the targeting reliability of EEG cap and the Beam F3 method. Further investigations by Cardenas^10^ and Kinjo^19^ et al. have examined the efficacy of targeting to the predictor of sgACC functional connectivity using the Beam F3 method. Beyond reliability and accuracy, additional factors such as speed, training efficiency for novices, and required external conditions hold significant value for clinical implementation. However, comprehensive research into the overall performance of these methods remains insufficient.

The present study aims to systematically compare the EEG cap, Beam F3, and CPC F3, and adjusted Beam F3 methods simultaneously, focusing on accuracy, reliability, speed, training efficiency and requirements. By providing comprehensive empirical evidence, this research explore which method offers a more accurate, consistent, and quicker approach to targeting DLPFC. This assists in clinical decision-making and helps advance towards a standardized clinical protocol, thereby enhancing the reliability and consistency of clinical findings.

## Methods

### 2.1 Operational procedures for each scalp-based targeting method

Prior to employing the four scalp-based targeting methods, the four critical reference points-namely, the nasion (NZ), inion (IZ), and left and right preauricular points (AL/AR) - were visually identified^20^. The detailed operational procedures and specific requirements for each method are outlined in Table 1.

**Table 1.** Detailed operational procedures and requirements for each scalp-based targeting method.

| Method | Operational procedures | Requirements |
| --- | --- | --- |
| EEG cap | (1) Put on the EEG cap.<br>(2) Ensure the cap's left-right symmetry by first aligning the cap's midline with the Nz and then positioning the Cz point on the EEG cap halfway between AL and AR.<br>(3) Adjust the cap's anterior-posterior position by placing the cap's Cz point halfway between Nz and Iz.<br>(4) Mark the DLPFC measurement according to the position of F3 on the cap. | EEG cap, tape measure |
| Beam F3<sup>12</sup> | (1) Put on a swimming cap.<br>(2) Measure a line passing through the AL and AR, record the length as L1, and mark the midpoint as Cz1.<br>(3) Measure a line passing through the Nz, Iz, and Cz1, record the length as L2, and mark the midpoint as Cz.<br>(4) Measure the head circumference and record it as L3.<br>(5) Once measurements L1, L2, and L3 are obtained, enter them into the software package (http://clinicalresearcher.org/eeg/). This software will provide two key values, X and Y.<br>(6) Mark a point along the head circumference that is X cm from the midline towards AL.<br>(7) From Cz, measure Y cm along the line running towards the point identified in step 6 and mark this as the DLPFC measurement. | Swimming cap, tape measure, website or software |
| CPC F3<sup>14</sup> | (1) Put on the swimming cap.<br>(2) Measure a line passing through the AL and AR, and mark | Swimming cap, tape measure, |
|  | the midpoint as Cz1.<br>(3) Measure a line passing through the Nz, Iz, and Cz1, and mark the midpoint as Cz.<br>(4) Along the midline, mark the point located at 27% of the distance from Nz.<br>(5) Measure a line that runs from AL, through the point established in the previous step, towards AR. Mark the point at 34% of the distance from AL as the DLPFC measurement. | calculator |
| Adjusted Beam F3<sup>16</sup> | Follow the same steps (1) to (6) as described for the Beam F3 method.<br>(7) From Cz, measure Y+0.35 cm along the line running towards the point identified in step 6 and mark this as the DLPFC measurement. | Swimming cap, tape measure, website or software |

### 2.2 Comparative Experiment

#### 2.2.1 Preparation

Ten individuals were selected from a 114-person MRI database (for details of MRI acquisition, please refer to 13). To enhance the generalizability of our findings, the selection strategy aimed to incorporate diversity in head size and sex. We selected five individuals with a head circumference ranging from 58 to 59 cm and another five with a head circumference ranging from 54 to 55 cm, of which six were men and four were women. Their mean and standard deviation (SD) of age was 19 ± 0.88 years. Virtual 3D polygonal mesh models of each individual’s head surface were generated based on the structural MRIs of the respective individuals (Fig. 1a), using the headreco command within SimNIBS software^21^. A 3D printing company subsequently produced ten 1:1 scale epoxy resin models with a precision of 50 μm, as illustrated in Fig. 1a. Prior to printing, conical holes were generated at each of the 10-20 system positions^22^, each with a radius of 1 mm and a depth of 2.6 mm, to facilitate coregistration with the MRI. The facial area from the nose to the mouth in the head models was anonymized, preserving the integrity of the head surface and the 10-20 positions.

**Figure 1.**
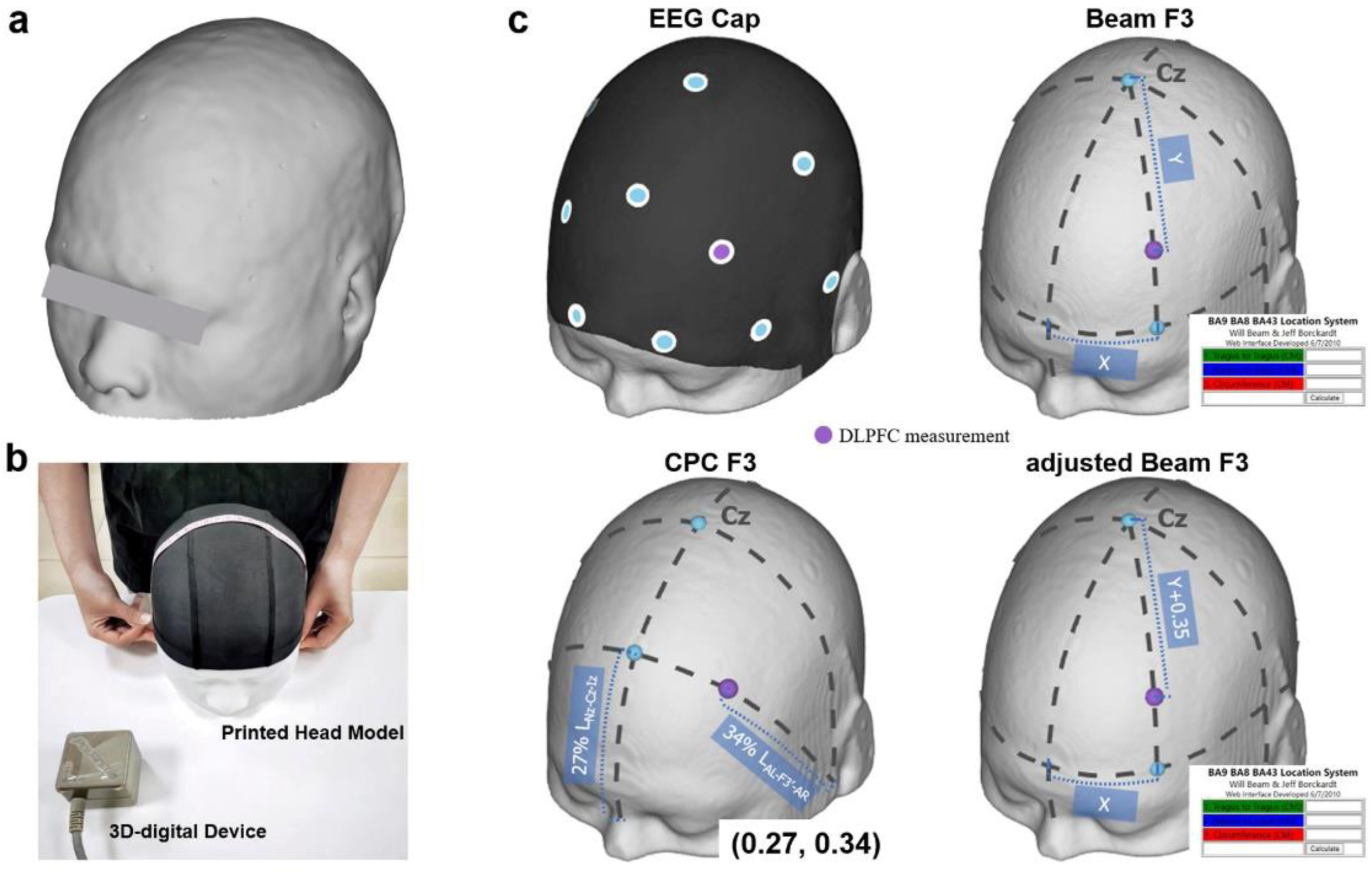
Comparative Experiment. **(a)** 3D printed head model of one selected individual. **(b)** The targeting setup includes a 3D printed head model wearing a black swimming cap, along with a 3D digitizer device for recording the coordinates of the measured points. **(c)** Diagrams illustrating the four compared methods.

#### 2.2.2 Experimental procedures

Three technicians, who were familiar with TMS, repeated each of the four scalp-based targeting methods twice on each of the 10 3D-printed head models, with an interval of at least 10 days between the repetitions (resulting in a total of 60 measurements). Prior to the experiment, they practiced these methods for approximately 10 hours in total to achieve stable performance. The targeting setup is shown in Fig. 1b. Using one of the four methods depicted in Fig. 1c, the technician performed the DLPFC measurement and the measurement time was recorded. Due to the nearly identical measurement steps of Beam F3 and adjusted Beam F3, both methods shared the same steps 1-6. The sequence in which the targeting methods - EEG cap, Beam F3, and CPC F3 - were applied was counterbalanced.

#### 2.2.3 Targeting performance Indices and Statistical Analysis

***Targeting accuracy*** of a method was quantified in two aspects. First, we defined the targeting error^23^ as the smallest perpendicular distance from the normal vector line at the DLPFC measurement site to the optimal group target. This target, identified as the sgACC anti-correlated peak^24^, is located at the Montreal Neurological Institute (MNI) coordinate [X-42 Y+44 Z+30]. Second, to assess targeting efficacy further, we caculated the electric field strength within the defined optimal group target area, a 12 mm sphere centered on the target^24^. We also examined a smaller, commonly used 5 mm^25^ and 10 mm^15^ sphere. Electric field simulations were conducted using simNIBS^21^, with the DLPFC measurement designated as the stimulation site and the stimulation direction oriented at 45 degrees from the anterior-posterior axis^26^; all other parameters were set to default. ***Targeting reliability*** was evaluated from two perspectives, in alignment with the definitions by Trapp^18^ and Jiang^14^: Inter-technician variability, which assesses method consistency against a group-averaged measurement coordinate derived from multiple technicians measuring the same individual, and intra-technician variability, which measures the consistency of measurements taken by the same technician on the same individual across two sessions. ***Speed*** of each method was determined by the time required to complete specific steps: Steps 1-4 in the EEG cap, Steps 1-7 in the Beam F3, and Steps 1-5 in the CPC F3. ***Training efficacy*** was assessed by the disparity in targeting error between three trained technicians, previously mentioned, and two newly recruited novice technicians who had no prior experience with any scalp-based targeting methods. The targeting procedure for novice technicians remained consistent with that described for trained technicians in section 2.2 (one session, 20 measurements in total). Prior to targeting, novices underwent approximately 30 minutes of study.

Statistical analyses were conducted as follows: A two-factor (method × session) repeated measures ANOVA was used for accuracy and time; reliability was analyzed using a repeated measures ANOVA; for training efficacy, a Wilcoxon rank-sum test compared the targeting errors between trained technicians and novices. Post hoc tests from the ANOVAs were conducted with Bonferroni corrections for multiple comparisons.

## Results

Measurements from four methods were projected onto the cortex (Fig. 2a). For the targeting error targeting the optimal group target, a significant main effect of method was observed (F(3,27) = 13.488, p < 0.001, Fig. 2b). The mean targeting errors (± SDs) in mm for each method were as follows: EEG cap = 7.67 ± 3.73, Beam F3 = 6.08 ± 4.06, CPC F3 = 4.16 ± 2.07, and adjusted Beam F3 = 6.93 ± 3.65. The targeting error for CPC F3 was significantly lower than those for the other methods (p < 0.05). No significant main effect of session (F(1,29) = 1.575, p = 0.220) or interactions were found (F(3,27) = 1.375, p = 0.256). Similarly, a significant main effect of method was found for the induced electric field strength at the optimal group target (F(3,27) = 5.167, p = 0.002, Fig. 2c). The mean electric field strengths (± SDs) in V/m for each method were as follows: EEG cap = 1.23 ± 0.13, Beam F3 = 1.25 ± 0.13, CPC F3 = 1.27 ± 0.11, adjusted Beam F3 = 1.26 ± 0.14. CPC F3 showed significantly stronger electric fields compared to EEG Cap (p < 0.001), with no significant differences noted with Beam F3 and adjusted Beam F3 (p < 0.05). Homogeneous results were observed for the electric field strength within the target, defined as both 5 mm and 10 mm spheres, indicating that the sphere size did not influence the outcomes.

**Figure 2.**
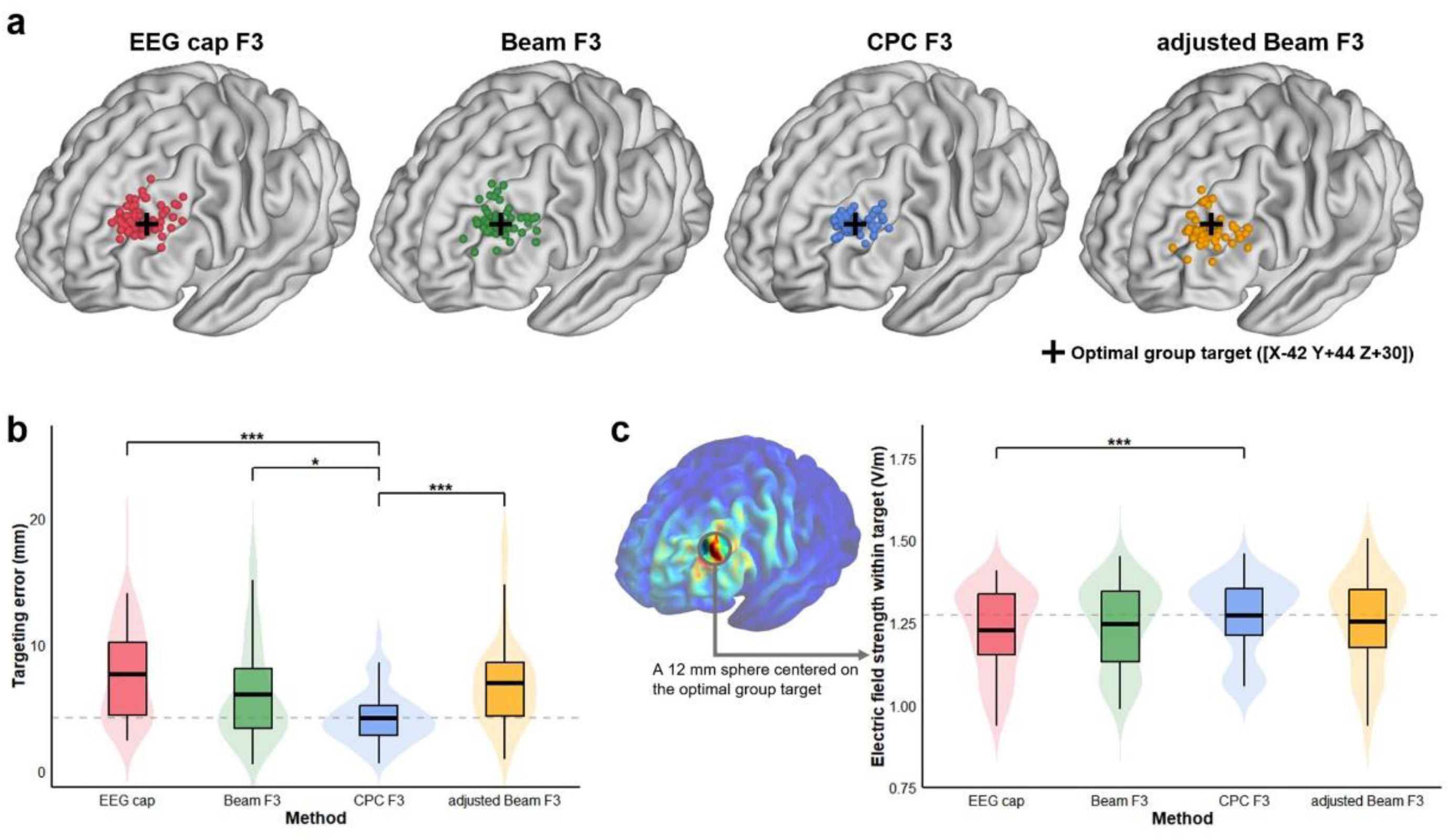
Comparison of targeting accuracy. **(a)** The scatter plot displays 60 measurements projected onto the MNI152 cortex for each method. A cross marks the optimal group target, noted for its strong normative anti-correlation with the sgACC. **(b)** Targeting error for the optimal group target. **(c)** Electric field strength within the optimal group target. A bold black line in the boxplot represents the mean performance across methods, while a dashed grey line denotes the mean for the CPC F3 method. Colors represent methods as follows: Red for EEG cap, Green for Beam F3, Blue for CPC F3, and Yellow for adjusted Beam F3. * indicates p < 0.05, ** indicates p < 0.01, *** indicates p < 0.001.

For reliability, significant differences were observed for both inter-(F(3,57) = 22.459, p < 0.001, Fig. 3a) and intra-technician variability (F(3,57) = 17.506, p < 0.001, Fig. 3b). The mean inter-technician variability (± SDs) in mm for each method were as follows: EEG cap = 6.40 ± 3.65, Beam F3 = 6.00 ± 3.58, CPC F3 = 3.30 ± 1.62, adjusted Beam F3 = 5.94 ± 3.60. Similarly, the mean intra-technician variability (± SDs) in mm for each method were as follows: EEG cap = 4.55 ±2.50, Beam F3=4.37 ± 2.53, CPC F3 = 2.32 ± 1.15, adjusted Beam F3 = 4.28 ± 2.53. Inter- and intra-technician variability for CPC F3 were significantly lower than those for the other three methods (p < 0.001).

**Figure 3.**
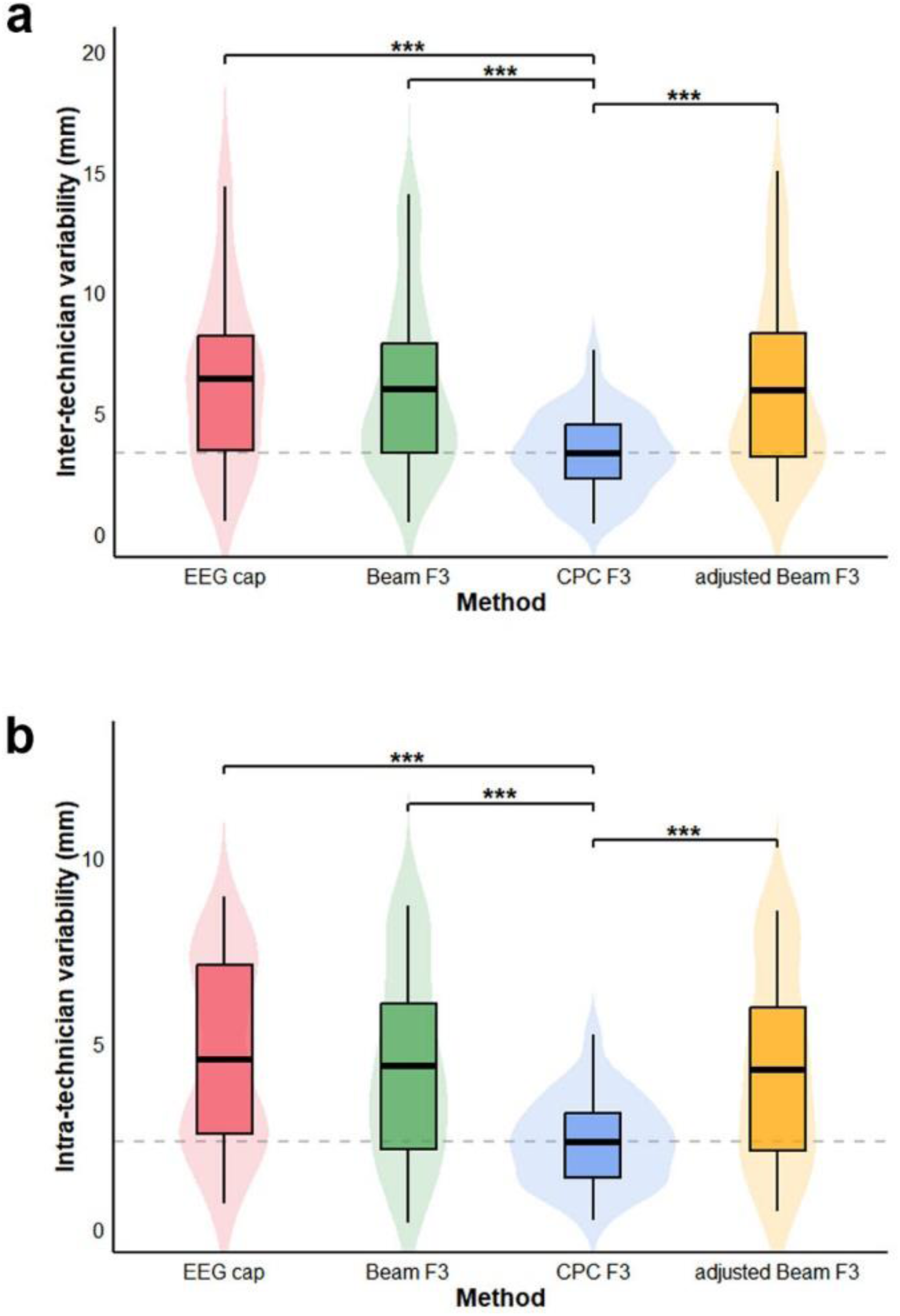
Comparison of targeting reliability. **(a)** Inter- and **(b)** intra-technician variability. A bold black line in the boxplots represents the mean variability across methods, while a dashed grey line denotes the mean for the CPC F3 method. ***: p-value<0.001.

Regarding speed, times for the adjusted Beam F3 were not recorded due to similar procedures with Beam F3. A significant main effect of the method was observed (F(2,28) = 98.510, p < 0.001). The mean time (± SDs) in seconds for each method were: EEG cap = 80 ± 2.7, Beam F3 = 140 ± 3.9, CPC F3 = 105 ± 3.1. The EEG cap was significantly faster than both CPC F3 and Beam F3 (p < 0.001), with CPC F3 being faster than Beam F3 (p < 0.001). A significant main effect of session (F(1,29) = 17.058, p < 0.001) and interaction between method and session (F(1,28) = 4.874, p = 0.011) were observed, indicating quicker measurements in session 2. However, the sequence of measurement times for the three methods remained consistent across both sessions, with the EEG Cap being the fastest, followed by CPC F3 and Beam F3 being the slowest; significant differences were confirmed between them (p < 0.001). In training efficacy, the targeting errors for novices using the Beam F3 were significantly higher than those of trained technicians (p = 0.0352) and marginally higher with the EEG cap (p = 0.0692). No significant differences were noted with CPC F3 or adjusted Beam F3 (p > 0.05).

To directly illustrate the performance distinctions among four scalp-based targeting methods, we collected evaluations on accuracy, inter- and intra-reliability, speed, training efficacy, and requirements, as shown in Fig. 4. The results showed that the radar plot for CPC F3 covers the largest area, suggesting it has the best comprehensive performance among the four methods.

**Figure 4.**
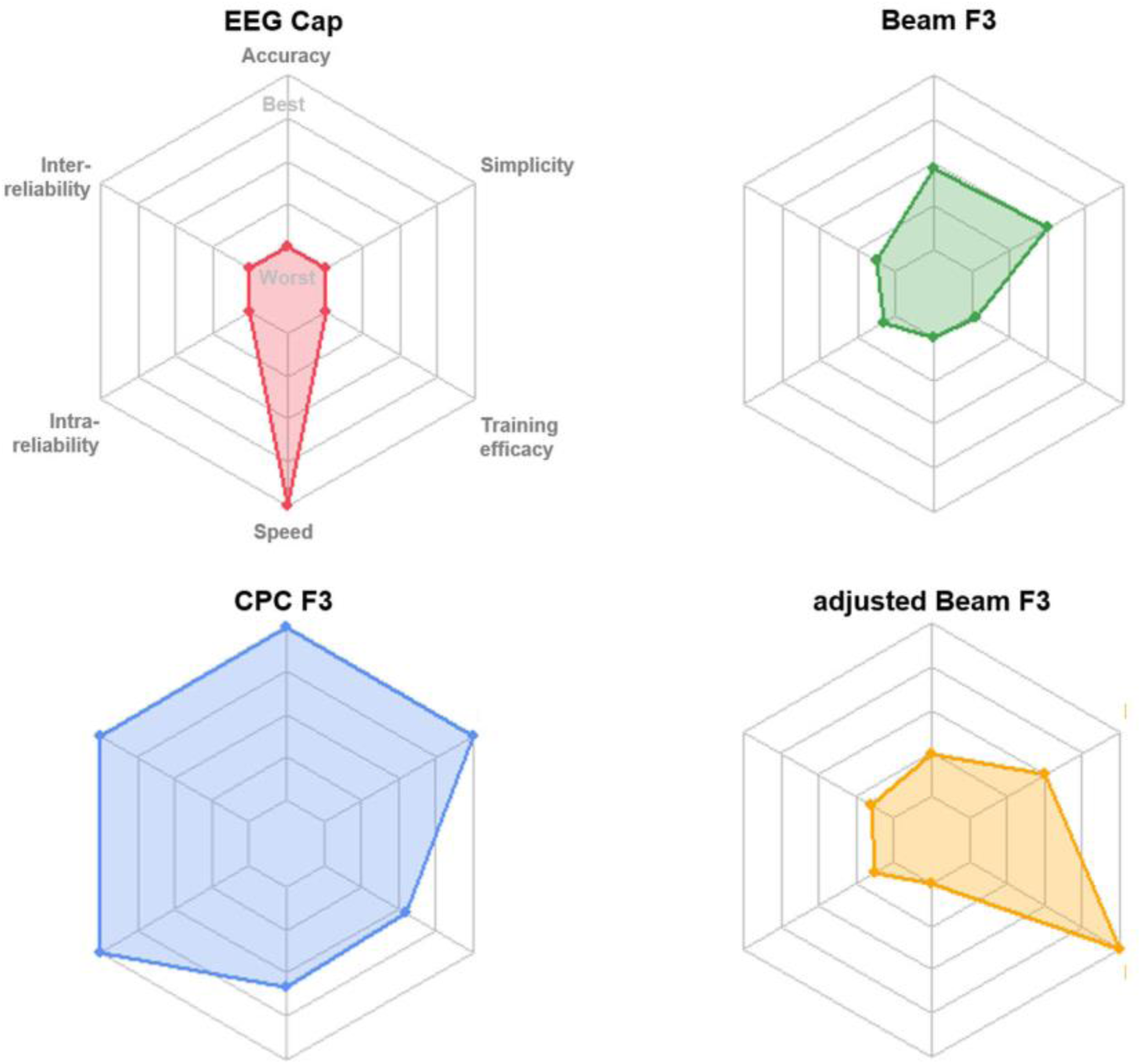
Comparison of targeting performance across six metrics. All metrics were transformed by negation and normalization from the mean value of each method, resulting in higher values representing better performance for each metric. Specifically, the accuracy metric was defined as the product of targeting errors and the reciprocal of electric field strength within the optimal group target. Training efficacy reflects the difference in mean targeting errors between trained technicians and novices. Simplicity indicates that the method requires fewer external conditions and is easier to implement.

## Discussions

Applying high-frequency pulses to the left DLPFC is the most commonly used protocol in the treatment of depression using rTMS^3^. Several scalp-based targeting methods - including the EEG cap, Beam F3, CPC F3, and adjusted Beam F3 - have been proposed to enhance the protocol implementation in clinical settings. Mastery of the scalp-based targeting methods has been recommended in rTMS practice guidelines for clinical practitioners of TMS^27^. In the present study, we systematically compared these methods for targeting the left DLPFC. The EEG cap method offers the advantages of being the fastest and imposing the least cognitive load during measurement. However, it does come with certain limitations. It necessitates the additional purchase of an EEG cap, which also introduces variability due to potential differences in manufacturing quality across brands. The standard sizing options available (small, medium, and large) may not always provide an ideal fit for individuals with varying head shapes and sizes. These factors may contribute to substantial variation in the locations of EEG cap measurements as reported across different studies, evidenced by differences in MNI coordinates such as [X-57 Y+41 Z+55]^28^ and [X-25 Y+50 Z+42]^29^. Our results revealed that even within the same study, the EEG cap still exhibited the lowest reliability, and inter-rater reliability values reported in previous studies were even worse ^17, 18^, approximately 9-10 mm.

The Beam F3 method offers advantages over the EEG cap method, as it does not require an additional EEG cap and can be manually measured in clinical settings by assessing geometric parameters on the scalp surface. Although the reliability of the Beam F3 method was found to be significantly higher than that of the EEG cap, the simplified targeting procedure of Beam F3 increases the cognitive load on technicians, particularly in recording the three measurement lengths (i.e., L1, L2, and L3). To avoid forgetting the records and the need to repeat measurements, technicians were required to input these records into software after each length measurement, which can be time-consuming. No statistically significant differences were found in accuracy between the Beam F3 and EEG cap methods, whether in targeting error or the electric field strength within the optimal target group. This finding is consistent with results from comparisons based on MRI scans, which did not involve measurement error. Cardenas et al.^10^ reported no significant differences in functional connectivity between the sgACC and the brain targets of the two methods, suggesting comparable targeting efficacy between Beam F3 and EEG Cap. In both methods, novices tended to generate significantly or marginally larger targeting errors than trained technicians, and they required additional training to achieve the same level of accuracy as trained technicians. Regarding the adjusted version of Beam F3, no significant improvement in accuracy was found. We attempted to change the coordinate of the optimal group target to [X-38 Y+44 Z+26] as suggested by Fox et al..^15^ Even though it was derived from a smaller sample size, this remains the target for the adjusted Beam F3^16^. However, no significant improvement was observed. We speculated that this might be due to the fixed shift (i.e., Y + 0.35 cm) not accommodating the variance in head sizes. Moreover, the nearly identical targeting procedures resulted in no significant differences in reliability or speed between Beam F3 and adjusted Beam F3, yet differences in training efficacy were noted. This may be related to the non-counterbalanced order of measurement between them, with the adjusted Beam F3 always following the Beam F3.

Surprisingly, CPC F3 exhibited remarkable targeting accuracy, achieving the lowest targeting error (4.16 mm) and significantly higher electric field strength within the target compared to the EEG Cap method. Our recent work^23^ presented the large-sample-based average CPC of the optimal group target (0.27, 0.33), which closely aligns with the average CPC for F3 (0.27, 0.34). We analyzed a 114-person MRI database to calculate the targeting errors for CPC F3 and CPC (0.27, 0.33) in targeting the optimal group target, revealing mean errors of 3.09 mm and 2.77 mm, respectively. The study by Cardenas et al.^10^ also reported no significant differences in the cortical area and network between the brain targets identified by neural navigation for the optimal group target and those identified by EEG F3, which is the target for CPC F3. These close targeting results explain the high accuracy observed with the CPC F3 method, even when targeting the optimal group target. Furthermore, no significant differences were found in the electric field strength within the optimal group target when comparing the Beam F3 and adjusted Beam F3 methods, suggesting comparable targeting efficacy among these methods. In addition to its accuracy, CPC F3 demonstrated practical advantages such as high reliability, moderate speed, and simple implementation, requiring only a swimming cap, tape, and a calculator. Our study also indicated that with only about 10 minutes of studying (30 minutes for four methods in total), a novice could achieve targeting accuracy comparable to that of trained technicians. The excellent targeting performance of the CPC F3 method could potentially be extended to the localization of other 10-20 points. For example, CPC (0.13, 0.78) for F8 targets the Inferior Frontal Gyrus for treating post-stroke aphasia^30^, and CPC (0.13, 0.78) for P3 (0.73, 0.34) targets the intraparietal sulcus for addressing post-stroke hemispatial neglect^31^.

We also investigated the impacts of head circumference on the accuracy of DLPFC targeting. No significant difference was observed in measurement precision between five individuals with large head sizes (58-59 cm) and five with small head sizes (54-55 cm), suggesting that accuracy is robust across different head sizes. Additionally, the difference between the 3D-printed head model and an actual head did not result in large differences in measurement time. We conducted an experiment in which three trained technicians performed the targeting procedure on newly recruited participants (15 measurements in total). The measurement times for the Beam F3 and CPC F3 methods showed no differences when performed on participants compared to the head model (p > 0.05, Wilcoxon rank-sum test). For the EEG cap method, a delay of 12 seconds was observed when conducted on participants compared to the head model (p = 0.048, Wilcoxon rank-sum test).

Regarding the limitations of our study, we used a 3D-printed head model to calculate targeting accuracy on MRI, which did not account for the influence of head movement and hair. Additionally, when using scalp-based targeting methods, there is potential for eye-hand coordination errors during the placement of the coil at the determined DLPFC position. However, this error is consistent across all four methods and should not impact our comparative conclusions. Notably, the iteration process for determining Cz in the CPC F3 method was simplified compared to the original definition provided by the tutorial video in Jiang^14^, aiming to improve comparability with other methods. The determination of Cz is a critical step in scalp-based targeting methods and warrants further investigation. Lastly, future studies may benefit from including larger and more diverse samples for a more comprehensive understanding.

## Conclusion

Our study provides a comprehensive evaluation and comparison of four scalp-based targeting methods for the DLPFC, focusing on key performance metrics such as targeting accuracy, reliability, speed, training efficacy, and implementation requirements. These insights can inform clinical decision-making and enhance the practical applicability of these methods. Notably, the CPC F3 method distinguishes itself as particularly promising, exhibiting superior accuracy, reliability, and training efficacy, alongside straightforward implementation and favorable speed characteristics, thereby underscoring its strong potential for clinical application.

## Acknowledgment

We thank Yilong Xu and Haosen Cai for their assistance in conducting the experiments.

## Author Contribution

Yihan Jiang: Conceptualization, Methodology, Data analysis, Investigation, Funding acquisition, Writing - original draft, review & editing, Visualization; Yuanyuan Chen: Conceptualization, Methodology, Data collection, Data analysis, Investigation, Writing - original draft, review & editing, Visualization; Lijiang Wei: Conceptualization, Data collection; Farui Liu: Methodology; Zeqing Zheng: Data collection; Zong Zhang: Methodology; Zheng Li: Writing - review & editing; Yingying Tang: Writing - review & editing; Jijun Wang: Writing - review & editing; Chaozhe Zhu: Conceptualization, Funding acquisition, Investigation, Project administration, Resources, Writing - review & editing.

## Statement

During the preparation of this work the authors used chatGPT in order to improve the readability and language of the manuscript. After using this tool, the authors reviewed and edited the content as needed and take full responsibility for the content of the published article.

